# Site-Specific Immune and Stromal Architecture Drive Resistance to Trastuzumab Deruxtecan in HER2+ Metastatic Breast Cancer

**DOI:** 10.1101/2025.05.23.654950

**Authors:** Glori Das, Matthew Vasquez, Jeffrey Zhang, Wei Yang, Yuan Gao, Xiaoxian Li, Hong Zhao, Stephen T.C. Wong

**Affiliations:** Department of Systems Medicine and Bioengineering, Houston Methodist Neal Cancer Center, Houston Methodist Hospital; Department of Biomedical Engineering, Texas A&M University; Advanced Cellular and Tissue Microscopy Shared Resource, Houston Methodist Neal Cancer Center, Houston Methodist Hospital; College of Arts and Sciences, Texas A&M University; Department of Pathology and Laboratory Medicine, Emory University School of Medicine

## Abstract

Trastuzumab deruxtecan (T-DXd) is an antibody-drug conjugate (ADC) that has demonstrated remarkable efficacy in HER2+ and HER2-low metastatic breast cancer (mBC). Yet, 40–50% of patients fail to respond, and mechanisms underlying resistance, particularly those involving the immune microenvironment, remain poorly understood. To address this gap, we conducted spatial proteomic profiling of metastatic lesions from 25 HER2+ mBC patients treated with T-DXd, using the Bruker GeoMX Digital Spatial Profiler (DSP) platform. Regions of interest (ROIs) were segmented by CD45 expression to distinguish immune-rich versus immune-poor compartments, and spatial marker expression was compared between responders (R) and non-responders (NR). Spearman correlation analysis was used to assess spatial coordination among HER2, fibronectin, and immune markers. Non-responders exhibited greater HER2 heterogeneity, lower CD45+ immune infiltration, and consistent upregulation of fibronectin and granzyme B. In both bone and brain metastases, NR tissues showed strong correlations between fibronectin and T cell markers, indicating spatial immune exclusion. In contrast, NR lymph node metastases displayed negative correlations between fibronectin and immune markers and uncoordinated B–T cell clustering, suggesting an alternative resistance mechanism rooted in immune disorganization rather than exclusion. Collectively, these results implicate HER2 heterogeneity, fibrotic immune exclusion, and immune suppression as key drivers of T-DXd resistance across metastatic sites. By integrating spatial proteomics with clinical outcomes, this study provides novel insights into the spatial immune landscape of ADC resistance and identifies actionable biomarkers for patient stratification and combination therapy design, laying a critical foundation for developing biomarker-guided therapies and personalized treatment strategies to overcome ADC resistance in metastatic breast cancer.

**Translational Significance:** T-DXd represents a major advance in antibody-drug conjugate (ADC) therapies to address HER2 heterogeneity in tumors through bystander killing. However, treatment options following resistance to T-DXd remain limited, leaving a critical unmet need. As new ADCs continue to emerge, resistance will likely remain a persistent challenge across tumor types. This work extends beyond T-DXd resistance—it provides a framework to address resistance mechanisms common to other ADCs. By investigating immune-mediated mechanisms of ADC resistance, this research identifies therapeutic targets that can be leveraged to develop new treatment strategies. By integrating spatial biology with clinical data, this study offers unprecedented insights into immune-tumor interactions within the resistant tumor microenvironment. This approach deepens mechanistic understanding and enables personalized treatment strategies guided by spatially resolved biomarkers. This novel combination of spatial mechanistic research with clinical data represents advancement toward overcoming ADC resistance and improving outcomes for patients with few therapeutic options.

## Introduction

Trastuzumab deruxtecan (T-DXd) is an antibody-drug conjugate (ADC) that has shown great success in treating HER2+ cancers. T-DXd was approved by the FDA to treat HER2-low metastatic breast cancer (mBC) in 2022 and approved for all HER2+ solid metastatic cancers in April 2024. T-DXd targets multiple biological mechanisms to overcome heterogeneity in HER2-low tumors. The trastuzumab antibody binds to the HER2 receptor, blocking HER2-mediated tumor growth. Trastuzumab also activates antibody-dependent cell-mediated cytotoxicity (ADCC) and phagocytosis (Petricevic et al., 2013; Shi et al., 2015), further destroying the tumor and improving chemosensitivity. Deruxtecan (DXd) is a highly cytotoxic topoisomerase I inhibitor. Trastuzumab and DXd are bound by a cleavable peptide linker. When cells endocytose T-DXd after HER2 binding, lysosomal enzymes cleave the linker, releasing DXd. DXd’s lipophilicity allows it to diffuse out of the target cell and diffuse into nearby off-target cells, including HER2-cells. This is known as the bystander killing effect. T-DXd’s especially potent bystander killing effect overcomes HER2 heterogeneity, making it highly effective against HER2-low tumors.

Clinical trials have shown unprecedented outcomes for mBC patients treated with T-DXd. DESTINY phase I-III trials show ∼50-60% objective response rate for T-DXd, including for heavily pre-treated patients--a major improvement over the 20-30% objective response observed with other chemotherapy regimens, including the HER2-targeted ADC trastuzumab emtansine (T-DM1) (Andre et al., 2024; Hurvitz et al., 2023; Modi et al., 2022). An especially exciting result is that T-DXd is efficacious against mBC brain metastases (Bartsch et al., 2024; Harbeck et al., 2024). T-DXd is now considered the preferred second-line therapy for HER2+ mBC.

While T-DXd’s success is groundbreaking, the fact that the remaining ∼40-50% of patients do not respond is a considerable cause for concern. Patients who show resistance to one ADC often fail subsequent ADC therapies, leaving patients with few to no treatment options (Huppert et al., 2025). Patient studies into causes of T-DXd resistance have been limited. In the phase 2 DAISY trial, ∼70% of HER2 overexpressing patients showed objective response; simultaneously, ∼30% of patients termed HER2-nonexpressing also showed objective response to T-DXd. This indicates that factors beyond target expression explain T-DXd resistance (Mosele et al., 2023). Moreover, out of six HER2-overexpressing T-DXd resistant patients, four showed intratumoral uptake of T-DXd, indicating that poor cellular uptake by itself is also insufficient to explain resistance. A study of adaptive resistance in ten paired pre-versus post-T-DXd patient measurements indicates upregulated and mutated DNA repair pathways in nine out of the ten patients post-treatment (Lee et al., 2024), indicating DXd targets DNA replication. However, pre-T-DXd treatment factors that predispose patients to resistance remain underexplored.

What also remains underexplored is the nature of the immune response in T-DXd resistant mBC. HER2 itself serves as a target for immune recognition for B and T cells, enabling response to immunotherapy. “Immune-hot” tumors—characterized by increased T-cell infiltration—tend to be more sensitive to treatment, especially when combined with immune checkpoint inhibitors (Wang et al., 2023). “Immune-cold” tumors, on the other hand, display less immune infiltration and are therefore more likely to survive. Aside from tumor-intrinsic mutations that promote resistance to immune-mediated filling, fibroblast-produced extracellular matrix barriers to immune infiltration can also promote an immune-cold phenotype. Metastatic breast cancer sites tend to be more immune-cold than primary tumor sites (Szekely et al., 2018). In HER2+ tumors, pathological complete response is associated with increased CD8+ T cell production of interferon gamma as well as improved infiltration of M1 macrophages, NK cells, B cells, and Th17 CD4+ T cells (Onkar et al., 2023). How these immune cells are distributed in T-DXd resistant vs. responsive cells has yet to be defined.

This study performed spatial profiling of the tumor-immune microenvironment of mBC to understand differences between T-DXd responders (R) and non-responders (NR) in order to gain insights into resistance mechanisms. We hypothesize that spatial immune exclusion and HER2 heterogeneity contribute to T-DXd resistance, and that spatial proteomic profiling, such as Bruker GeoMX spatial proteomics, can reveal biomarkers predictive of response.

## Methods

### Clinical Samples

Metastasis tissue sections were obtained from 25 HER2+ mBC patients, aged 41-84 years, who had developed resistance to multiple chemotherapy regimens at Emory University Hospital. Following specimen collection, all patients received T-DXd treatment. Based on treatment outcomes, patients were classified as either responders or non-responders. All procedures were approved by the Houston Methodist Institutional Review Board, and informed consent was obtained from all patients.

Formalin-fixed, paraffin-embedded (FFPE) tissue slides were prepared from biopsies or surgical resections of mBC lesions from various organs. A total of 29 specimens were collected from 23 patients, with approximately half (n = 12) derived from T-DXd-responsive patients and the remaining (n = 17) from T-DXd-nonresponsive patients.

### Spatial Proteomics for Immune Profiling

The Nanostring GeoMX spatial proteomics platform utilizes antibodies connected with a UV-cleavable linker to fluorescent nucleotide barcodes, indicating both immune marker and spatial location. Tissues were treated with these nucleotide-barcoded antibodies from the NanoString Immune Cell Profiling Core Panel and the Immune Cell Typing Panel. Some of these antibodies are used as “morphology markers” via fluorescence imaging. In this case, the morphology markers were antibodies directed against PanCK (cancer), CD45 (white blood cells), and DAPI (cell nuclei). Fluorescence images were obtained and used for region of interest (ROI) selection. ROIs were selected from PanCK-high and PanCK-low regions. Six to eight ROIs were collected per tissue, with each ROI having a diameter of 300µm—coincidentally, the same diameter of a breast cancer biopsy needle (Fig. 3B, C). Within each ROI, CD45+ cells were segmented using the GeoMX software. In the segmented areas, UV light exposure detaches the nucleotide barcodes from the antibodies. These barcodes are aspirated from the sample and counted with NanoString’s nCounter Pro machine (Fig. 2). Nucleotide counts are used to determine immune marker expression within each ROI. Following QC and geometric mean normalization, expression data is studied via linear mixed modeling, with patient as a random effect since multiple ROIs come from the same patient.

**Figure 1:**
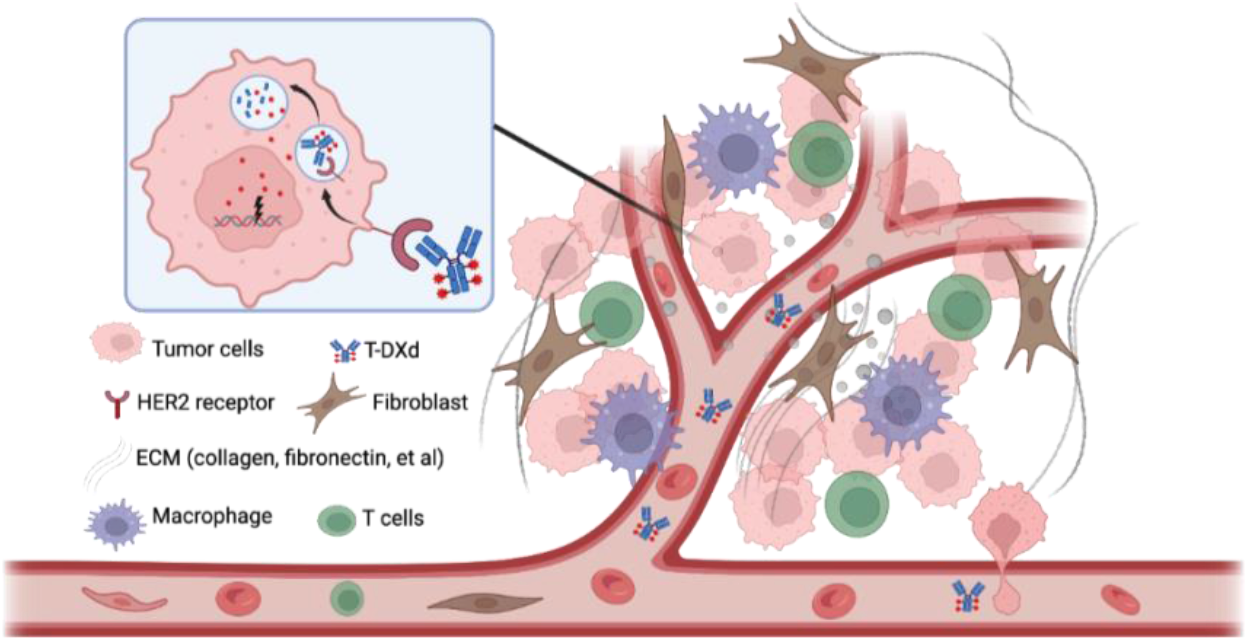
Tumor-immune microenvironment in T-DXd–treated tumors.

**Figure 2:**
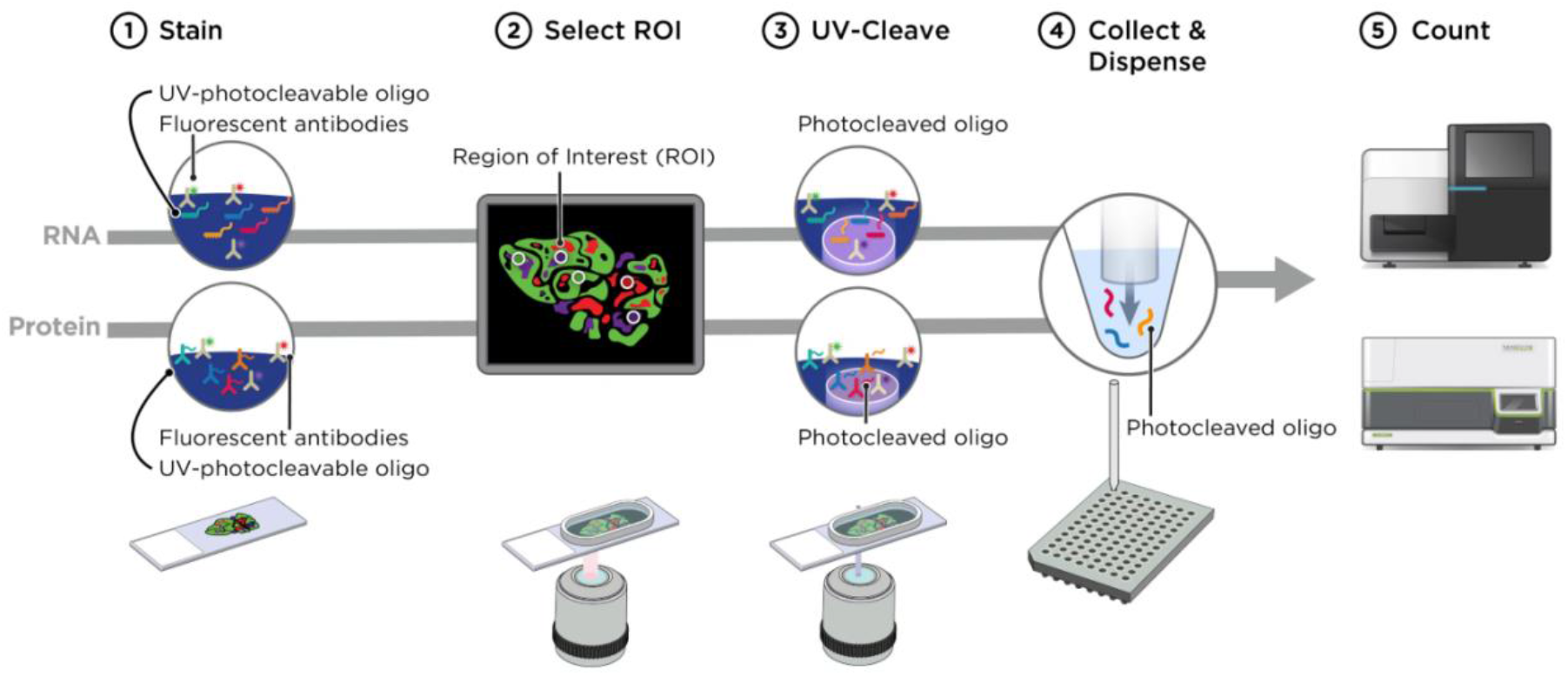
Nanostring GeoMX spatial profiling workflow. https://www.biochain.com/nanostring-geomx-digital-spatial-profiling/

**Figure 3:**
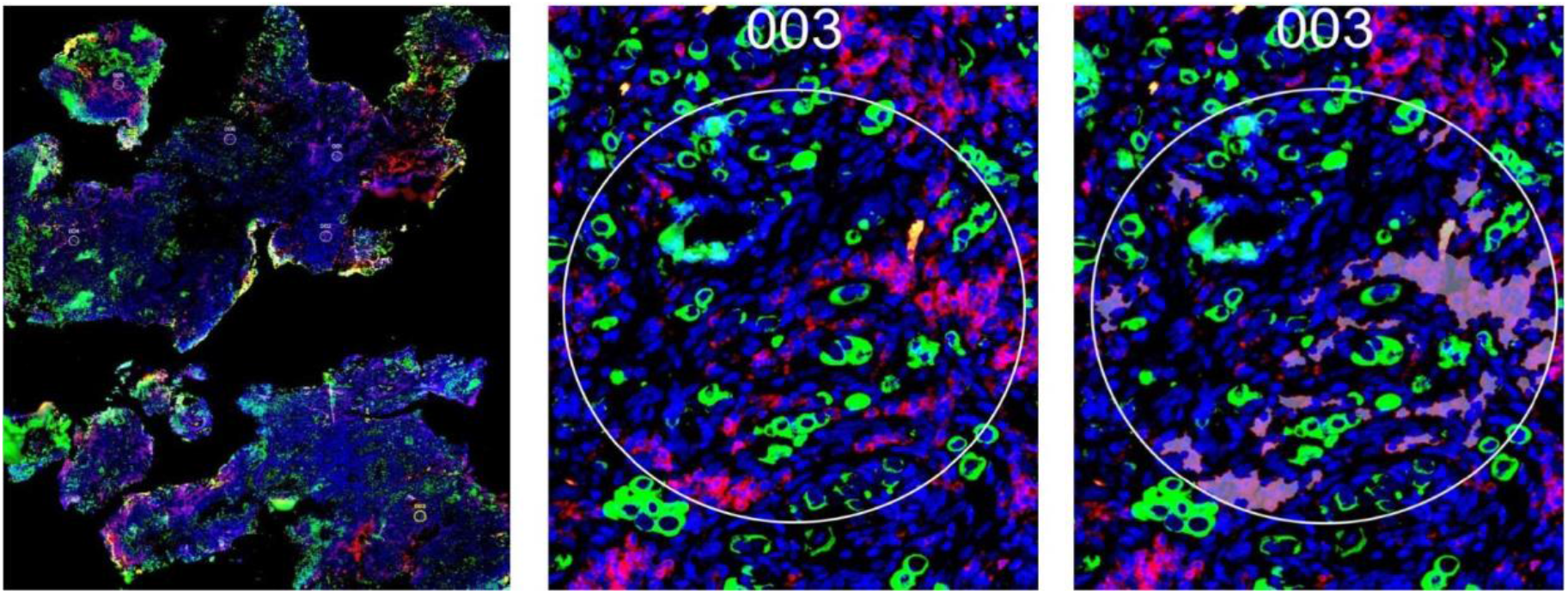
ROI selection and immune cell segmentation for oligonucleotide barcode collection. (A) Whole slide image with circled ROIs. Green = PanCK (cancer). Red = CD45 (immune). Blue = DAPI (nuclei). (B) Single ROI. Diameter = 300µm. N = 192. (C) Immune cells segmented for spatial profiling.

### HER2 Fluorescent Staining and Scoring of ROIs

In clinical practice, HER2 heterogeneity is typically overlooked when scoring biopsy samples. An overall HER2 score is assigned to the slides based on the intensity and area of brightfield HER2 staining. However, as mentioned before, HER2 heterogeneity plays a critical role in chemotherapy resistance and warrants closer examination. To capture spatial variation more precisely, ROIs were individually scored for HER2 expression and analyzed for corresponding local immune profiles across different HER2 levels, rather than relying solely on overall tissue scores.

ROIs were chosen without prior knowledge of their HER2 scores to avoid bias. Tissue sections adjacent to the ones used for GeoMX analysis were stained with fluorescently labeled anti-HER2 antibodies. Background subtraction of the fluorescent images was performed using CellSens (Evident Corporation, Tokyo, Japan) to optimize signal clarity. Fluorescence intensity was divided into quartiles for scoring HER2 expression. Using the image analysis software QuPath [ref], individual cells within each ROI were assigned scores of 0, 1+, 2+, or 3+, based on the quartile of HER2 intensity detected per cell (Fig 4).

**Figure 4:**
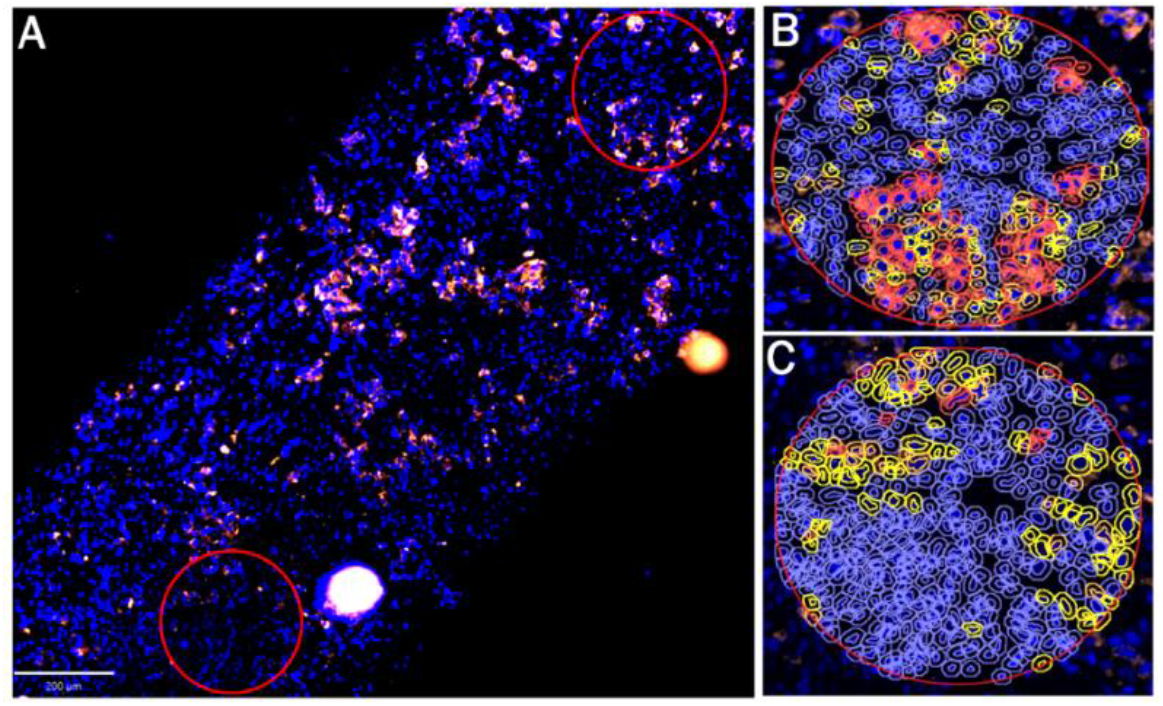
HER2 scoring by QuPath. (A) Fluorescently stained HER2 ROIs (red circles). (B) HER2 3+ ROI (top right of A): red = individual cell score of 3+, orange = 2+, yellow = 1+, blue = 0. (C) HER2 1+ ROI (bottom left of A).

Table 3 shows the methodology for HER2 scoring of ROIs, akin to brightfield scoring methods (Ivanova et al., 2024). Comparing QuPath scores to brightfield HER2 staining confirmed the validity of this method for ROI analysis.

**Table 1:** Patient demographics and treatment histories prior to specimen collection (n=25).

| Race | # Patients |
| --- | --- |
| Caucasian | 12 (48%) |
| African-American | 9 (36%) |
| Asian-American | 3 (12%) |
| Unknown | 1 (4%) |
| **Age** |  |
| 40s | 3 (13.0%) |
| 50s | 11 (47.8%) |
| 60s | 5 (21.7%) |
| 70s-90s | 5 (21.7%) |
| **Treatment Category** | **# Patients** |
| **Anti-HER2 therapies** (i.e. trastuzumab, pertuzumab) | 15 (65.2%) |
| T-DM1 | 10 (43.5%) |
| THP (docetaxel + pertuzumab + trastuzumab) | 8 (34.8%) |
| TCHP (THP + carboplatin) | 2 (8.7%) |
| Anti-HER2 + tyrosine kinase inhibitors | 3 (13.0%) |
| **Estrogen signaling inhibitors** (i.e. selective estrogen receptor modulators, aromatase inhibitors) | 15 (65.2%) |
| **Microtubule Targeting Agents** | 15 (65.2%) |
| **AC (doxorubicin + cyclophosphamide)** | 11 (47.8%) |
| **Palbociclib (CDK4/6 inhibitor)** | 7 (30.4%) |

**Table 2:** Organs represented in metastasis samples (n=29). R = response; NR = non-response.

| **Both R and NR** | **R Only** | **NR Only** |
| --- | --- | --- |
| Bone (7R vs. 4NR) | Pericardium (1) | Liver (3) |
| Brain (2R vs. 2NR) | Abdominal cavity (1) | Skin (1) |
| Lymph Nodes (2R vs. 2NR) |  |  |
| Chest Wall (2R vs. 2NR) |  |  |

**Table 3:** HER2 scoring of ROIs based on fluorescent HER2 staining and QuPath image analysis.

| HER2 Score for ROI | Scoring Method |
| --- | --- |
| 0 | <10% of cells in the ROI fall within the 2<sup>nd</sup>-4<sup>th</sup> quartiles of HER2 fluorescence brightness values (intensity) for the given slide. |
| 1+ | >10% of cells in ROI fall within the 2<sup>nd</sup> quartile of HER2 brightness values for the given slide, with <10% of cells in the higher quartiles. |
| 2+ | >10% of cells in ROI fall within the 3<sup>rd</sup> quartile of HER2 brightness values for the given slide, with <10% of cells in the higher quartiles. |
| 3+ | >10% of cells in ROI fall within the 4<sup>th</sup> quartile of HER2 brightness values for the given slide. |

**Table 4:** List of immune microenvironment markers chosen for GeoMx spatial proteomics.

| Set of Markers | GeoMx® Immuno-Oncology Kit | Proteomics Markers |
| --- | --- | --- |
| Core Markers | Immune Cell Profiling Core Panel | CD11c |
|  |  | PDL1 |
|  |  | PD1 |
|  |  | CTLA4 |
|  |  | Ki67 |
|  |  | CD3 |
|  |  | Granzyme B (GZMB) |
|  |  | CD56 |
|  |  | Smooth Muscle Actin (SMA) |
|  |  | Fibronectin |
|  |  | CD45 |
|  |  | CD68 |
|  |  | Beta-2-Microglobulin (B2M) |
|  |  | CD20 |
|  |  | CD8 |
|  |  | CD4 |
| Expanded Markers | Immune Cell Profiling Core Panel plus Immune Cell Typing Panel | All Core Markers |
|  |  | Fibroblast Activation Protein alpha (FAPα) |
|  |  | FOXP3 |
|  |  | CD163 |
|  |  | CD66b |
|  |  | CD14 |
|  |  | CD45RO |
|  |  | CD34 |

## Results

### Pan-Cancer Analysis Identifies Divergent Relationships Between HER2 Heterogeneity and Immune Infiltration Across Response Groups

We assessed differences in HER2 scores between responders and non-responders and examined associations between HER2 scores and immune infiltration. Whole-slide HER2 scoring confirmed that higher HER2 score was significantly associated with improved treatment response, as expected (Fig 5A). ROI-level analysis based on QuPath HER2 scores indicates that NR samples exhibited a greater proportion of ROIs with HER2 score 0 and showed higher variability in HER2 scores (Fig 5B). Consistently, a t-test comparing the standard deviation of HER2 scores across response groups revealed significantly greater heterogeneity in NR ROIs (t = 2.75, p = 0.013). To assess how relationships between HER2 score and immune infiltration differed between R vs. NR, correlation coefficients were compared using the Fisher r-to-z transformation. Surprisingly, markers suggestive of killer T cell activity showed negative correlations with increasing HER2 score in the response group, versus positive correlations existing in the NR group (Fig 5C).

**Figure 5:**
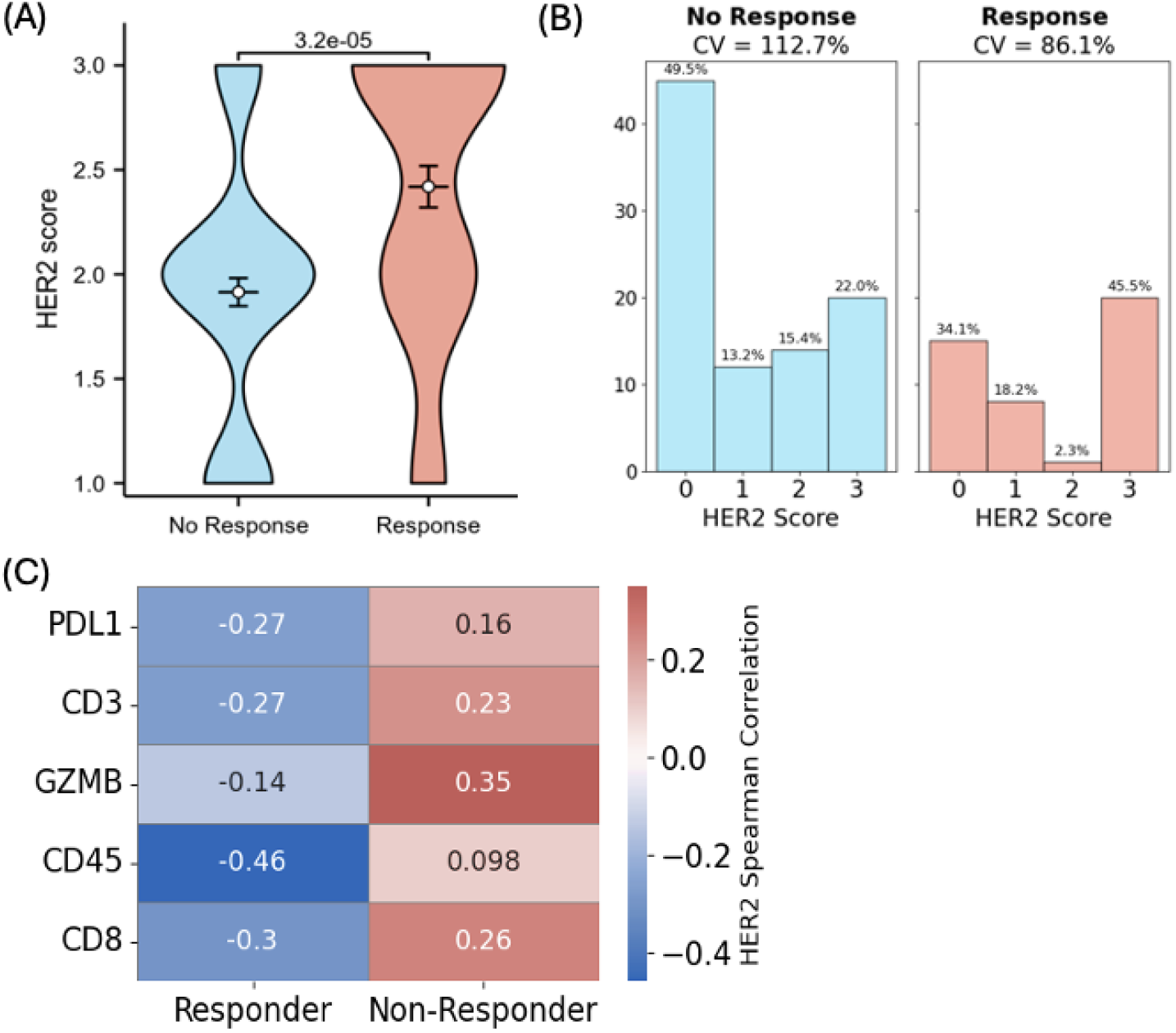
Relationship between HER2 heterogeneity, immune infiltration, and T-DXd response. (A) Distribution of whole-slide tissue scores by Emory pathologists (n = 12R, 17NR). (B) Distribution of HER2 scores for individual ROIs based on QuPath scoring (n = 7R, 14NR). CV = coefficient of variation. (C) Immune marker correlations with HER2 score between R vs. NR. P < 0.05 for all based on Fisher r-to-z transformation’ Cohen’s d ∼0.5 for all.

### Pan-Cancer Analysis Reveals Differential Immune Infiltration Patterns Between T-DXd Responders and Non-Responders at the ROI level

Differences in immune infiltration between R and NR were analyzed using a linear mixed-effects model, incorporating patient as a random effect to account for the non-independence of multiple ROIs per patient. A group of 17 core markers—derived from the Bruker Immune Cell Profiling Core Panel—was quantified across all tissue samples. To capture a broader spectrum of immune cell populations, a subset of samples was further studied with the Bruker Immune Cell Typing Panel, which included seven additional immune markers. These are referred to as the expanded marker set throughout the study. This approach enabled robust assessment of both core and extended immune signatures across the cohort, while controlling for within-patient variability.

Our linear mixed-effects modeling identified small but significant changes in immune and stromal marker expression, providing mechanistic insights into factors that may contribute to tumor resistance to T-DXd therapy. Core marker analysis showed that fibronectin was significantly upregulated in NR tissues, whereas CD45 was significantly upregulated in R tissues, suggesting fibronectin as a potential physical barrier to immune infiltration (Fig 6A). Natural killer (NK) cells (CD56) and dendritic cells (CD11c) were also elevated in R tissues. Conversely, GZMB (produced by T cells and NK cells), beta-2-microglobulin (B2M, part of the MHC I complex), and CD68 (macrophages) were elevated in NR tissues.

**Figure 6:**
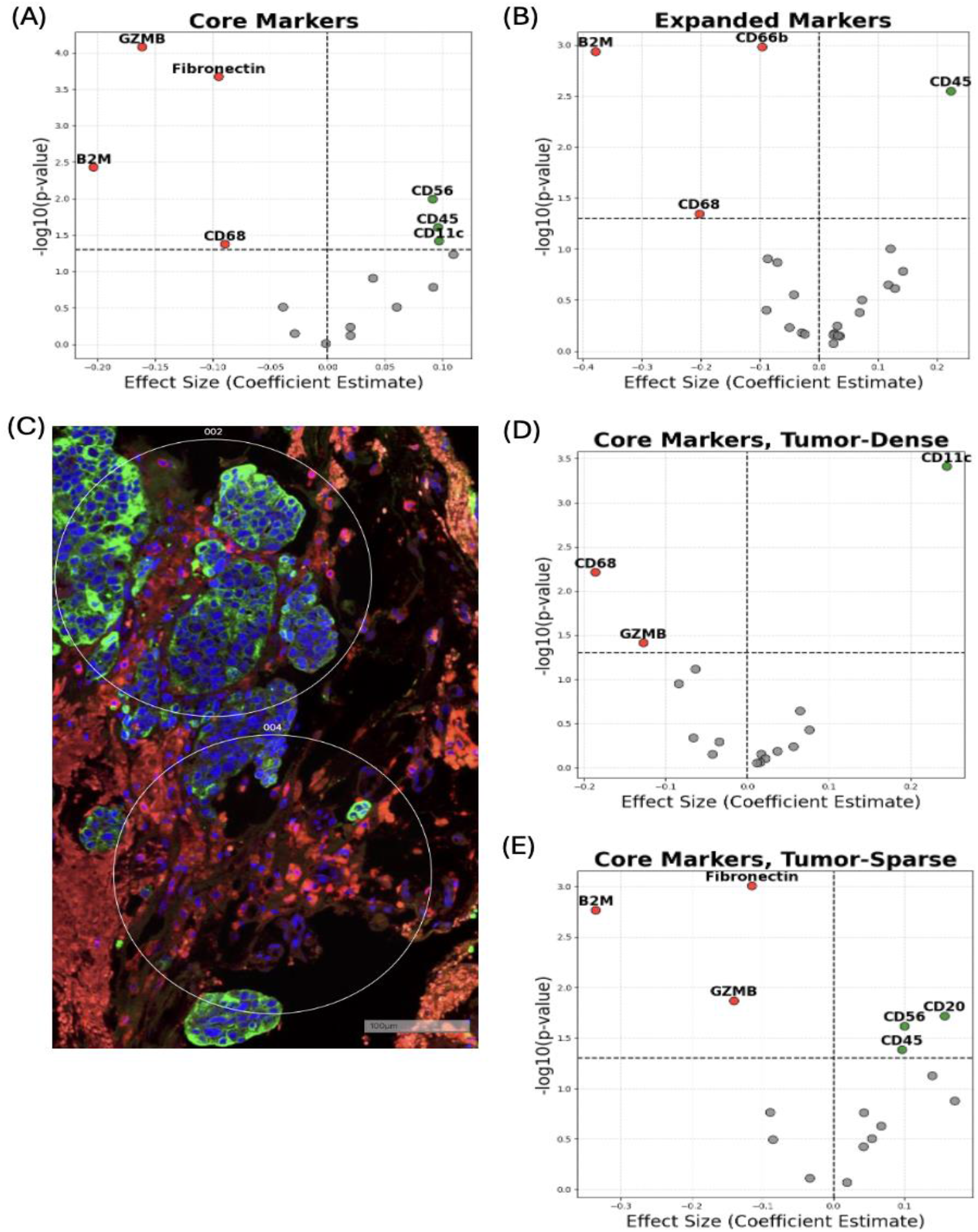
Differential immune infiltration in T-DXd R vs. NR pan-cancer tissues. (A) Core marker linear mixed model significant results (n = 12R, 17NR). GZMB = granzyme B, B2M = Beta-2-microglobulin. (B) Expanded marker linear mixed model significant results (n = 7R, 7NR). (C) Representative figure showing adjacent tumor-dense and tumor-sparse ROIs. (D) Tumor-dense ROI core marker linear mixed model significant results (n = 12R, 17NR). (E) Tumor-sparse ROI core marker linear mixed model significant results (n = 12R, 17NR). Red = higher in NR; Green = higher in R.

Analysis was done on the expanded marker set to assess whether these core findings were consistent. CD45 remained upregulated in R tissues, while B2M and CD68 persisted in being elevated in NR tissues. Additionally, CD66b (neutrophils) emerged as significantly upregulated in NR tissues.

ROIs were subsequently stratified as “tumor-dense” or “tumor-sparse” based on the mean number of PanCK+ cells within each organ. ROIs exceeding the organ-specific PanCK+ mean were labeled “tumor-dense,” and those below the mean were labeled “tumor-sparse” (see Fig. 6C for representative examples). Tumor-dense NR ROIs showed persistent upregulation of GZMB and CD68, whereas tumor-dense R ROIs showed persistent upregulation of CD11c (Fig 6D). Tumor-sparse NR ROIs showed persistent upregulation of GZMB, B2M, and fibronectin; tumor-dense R ROIs showed persistent upregulation of CD45, and CD56, as well as upregulation of CD20 (B cells) (Fig 6E).

To summarize, immune cell infiltration was consistently higher in R samples, especially in tumor-sparse regions. Elevated infiltration of CD11c, CD56, and CD20 (tumor-sparse ROIs) was associated with favorable response to T-DXd therapy. In contrast, fibronectin, especially in tumor-sparse regions, may contribute to therapeutic resistance by impeding immune cell infiltration, although macrophages and GZMB-producing cells remained infiltrative in NR samples.

### Pan-Cancer Analysis Indicates Roles of Fibronectin and GZMB in Predicting Response to T-DXd Based on Treatment History

GZMB and fibronectin consistently displayed small but significant changes negatively correlated with response. These markers were uniformly upregulated for NR among patients taking estrogen inhibitors, microtubule inhibitors, Herceptin-based therapies, Ibrance, and AC. This further affirms the concept of elevated GZMB and fibronectin as indicators of NR in pre-T-DXd tissues.

### Organ-Specific Analysis is Essential for Understanding the Immune Microenvironment

While pan-cancer analysis is helpful for providing general insights, the reality is that immune microenvironments differ between organs, which will create inter-organ differences in resistance mechanisms. Cluster analysis reveals that despite R vs. NR status, liver and brain samples still clustered together because of common immune microenvironment features—brain tissues in particular demonstrated a unique shared phenotype (Fig 7). Hence, the figures from now on will focus on organ-specific analysis.

**Figure 7:**
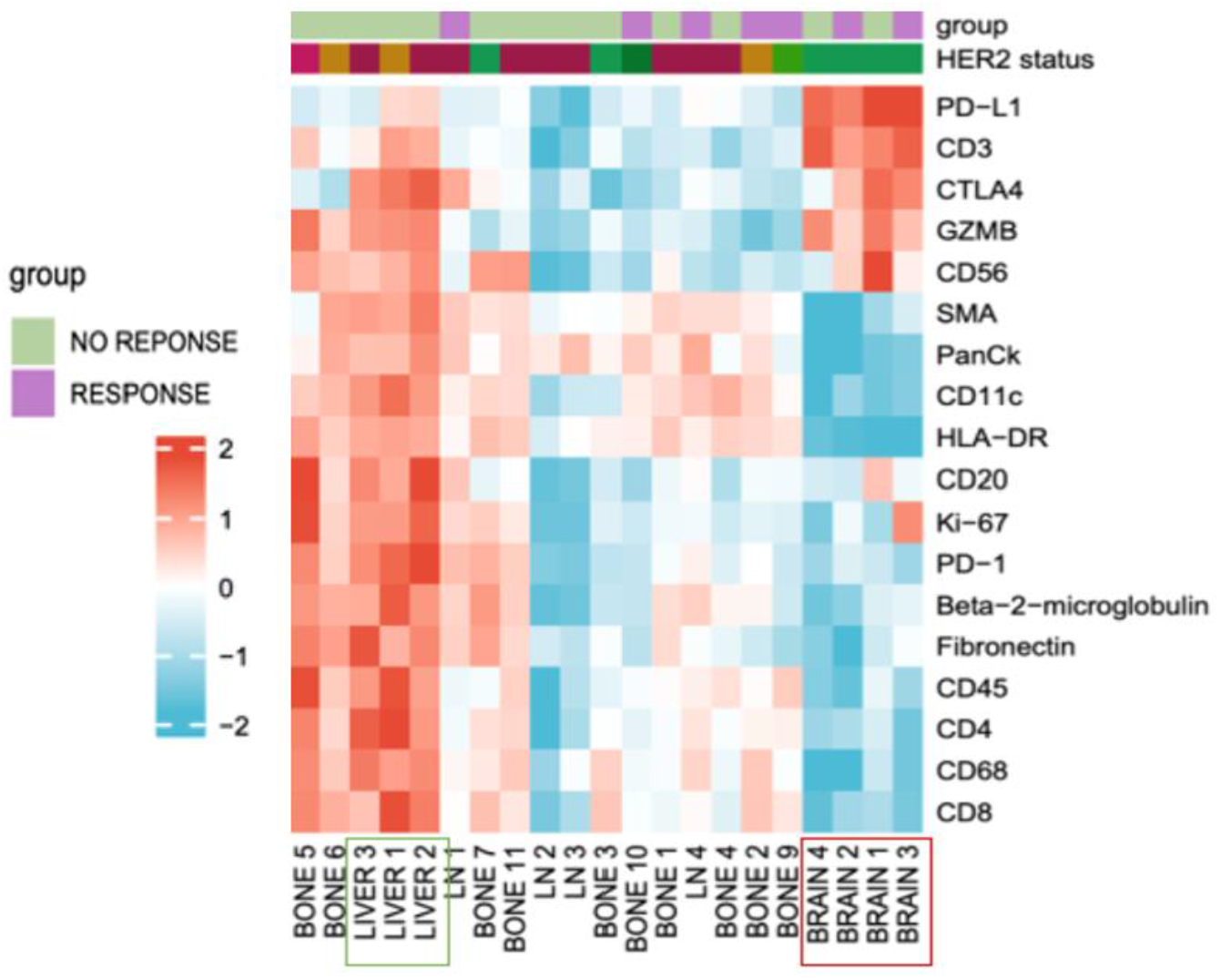
Organ clustering based on immune microenvironment factors.

### Lymph Node NR Metastases Feature Both Immunosuppression and Decreased Immune-Fibronectin Interaction

Cluster analysis of lymph node metastases revealed a marked distinction between responder (R) and non-responder (NR) profiles. Both immune and stromal markers were broadly downregulated in NR samples (Fig. 8A), suggesting a globally suppressed tumor microenvironment. Pairwise Spearman correlation analysis also indicates a stark difference in coordination of the immune microenvironment response between R vs. NR lymph nodes (Fig 8B). In R ROIs, strong positive correlations were observed between PDL1 and CD45 (ρ = 0.88), PDL1 and CD68 (ρ = 0.94), and PDL1 and CD4 (ρ = 0.88). Additionally, a robust correlation between SMA and Fibronectin (ρ = 0.92) was observed only in responders, indicating that extracellular matrix (ECM) remodeling is better coordinated in this group. The most striking observation is that NR demonstrated several significant negative correlations between Fibronectin and CD45 (ρ = – 0.24), SMA (ρ = –0.10), and CD56 (ρ = –0.12), suggesting exclusion of immune cells in regions of dense ECM. Furthermore, NR showed stronger internal coordination among lymphoid markers, including high correlations between CD20 and CD3 (ρ = 0.85), CD20 and CD8 (ρ = 0.89), and CD20 and CD4 (ρ = 0.96). These findings may reflect B–T cell clustering that is not associated with functional immune activation. Together, these findings suggest that responders exhibit more coordinated interactions between immune checkpoints, infiltrating immune cells, and ECM remodeling. In contrast, non-responders display patterns consistent with immune exclusion and potentially unproductive lymphoid clustering.

**Figure 8:**
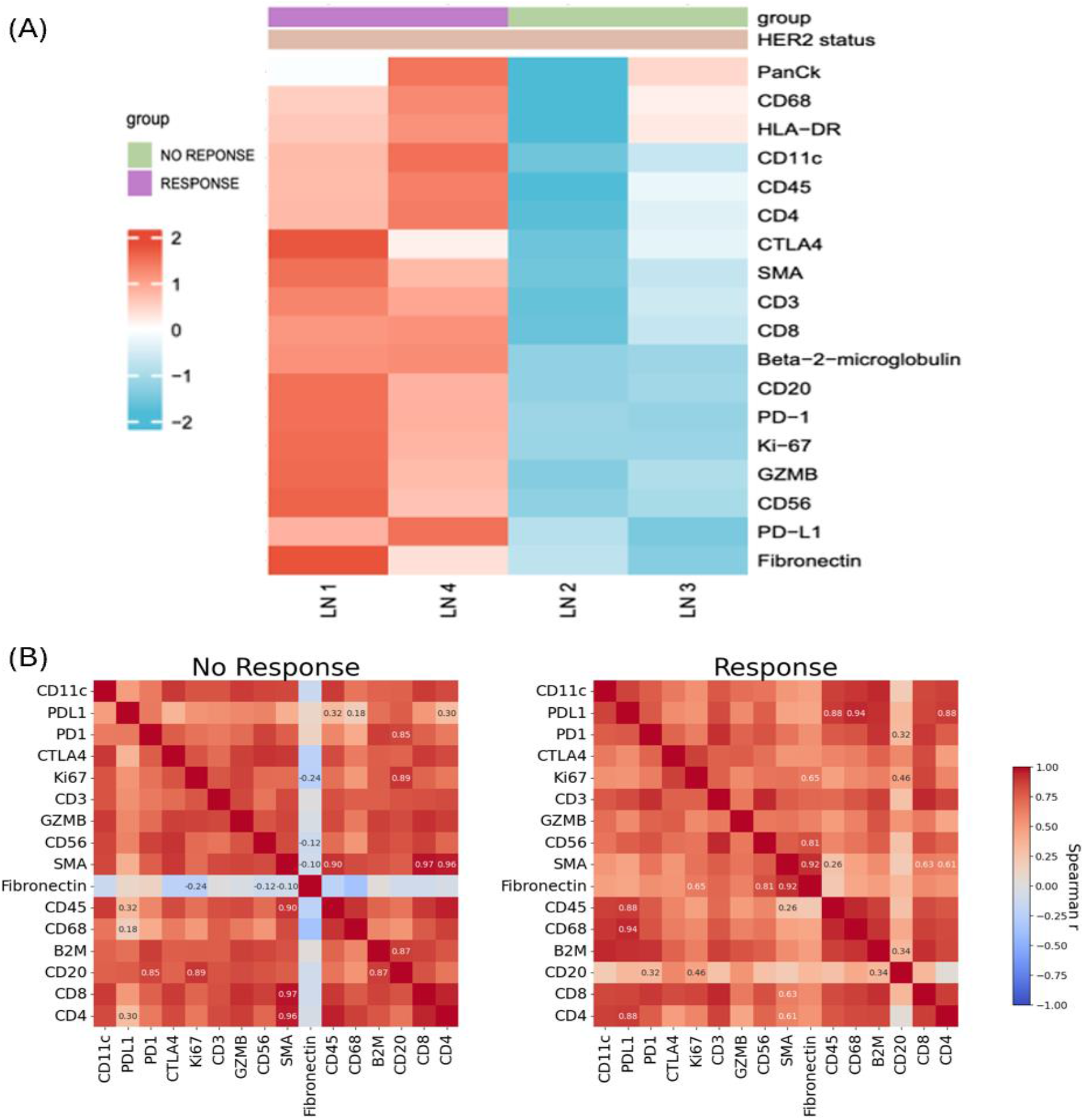
Distinct immune and stromal coordination patterns in lymph node R vs NR metastases. (A) Z-scored expression heatmap of immune and stromal markers in lymph node metastases, stratified by T-DXd response. (B) Spearman correlation matrices comparing marker–marker relationships in NR (left) and responder (R, right) lymph node samples. Only marker pairs with significant differences in correlation between groups (p < 0.05, Fisher z-test) are annotated.

### Fibronectin and Immune Marker Correlations Are Conserved Across Metastatic Bone and Brain Tumors in NR

Across both bone and brain metastases, NR tissues exhibited strong positive correlations between fibronectin and T cell markers, including CD3, CD4, and CD8 (Fig. 9A–B). In bone metastases, fibronectin expression correlated strongly with CD3, CD4, and CD8 (Spearman’s ρ > 0.79 for all), indicating tight spatial co-localization of fibrotic extracellular matrix (ECM) and T cell populations. A similar pattern emerged in brain metastases, with correlations ranging from ρ = 0.75 to 0.80 for the same markers. In contrast, these associations were markedly attenuated or even reversed in responders, for example, CD8 (ρ = –0.20 in brain; ρ = 0.33 in bone), suggesting that immune cells are spatially decoupled from fibronectin-rich ECM in responsive tumors. This conserved signature across bone and brain metastases implicates fibronectin as a potential spatial barrier to effective T cell infiltration in NR. Notably, this trend contrasts with lymph node metastases, where fibronectin-immune correlations followed an opposite pattern, underscoring site-specific differences in tumor–immune–ECM interactions.

**Figure 9:**
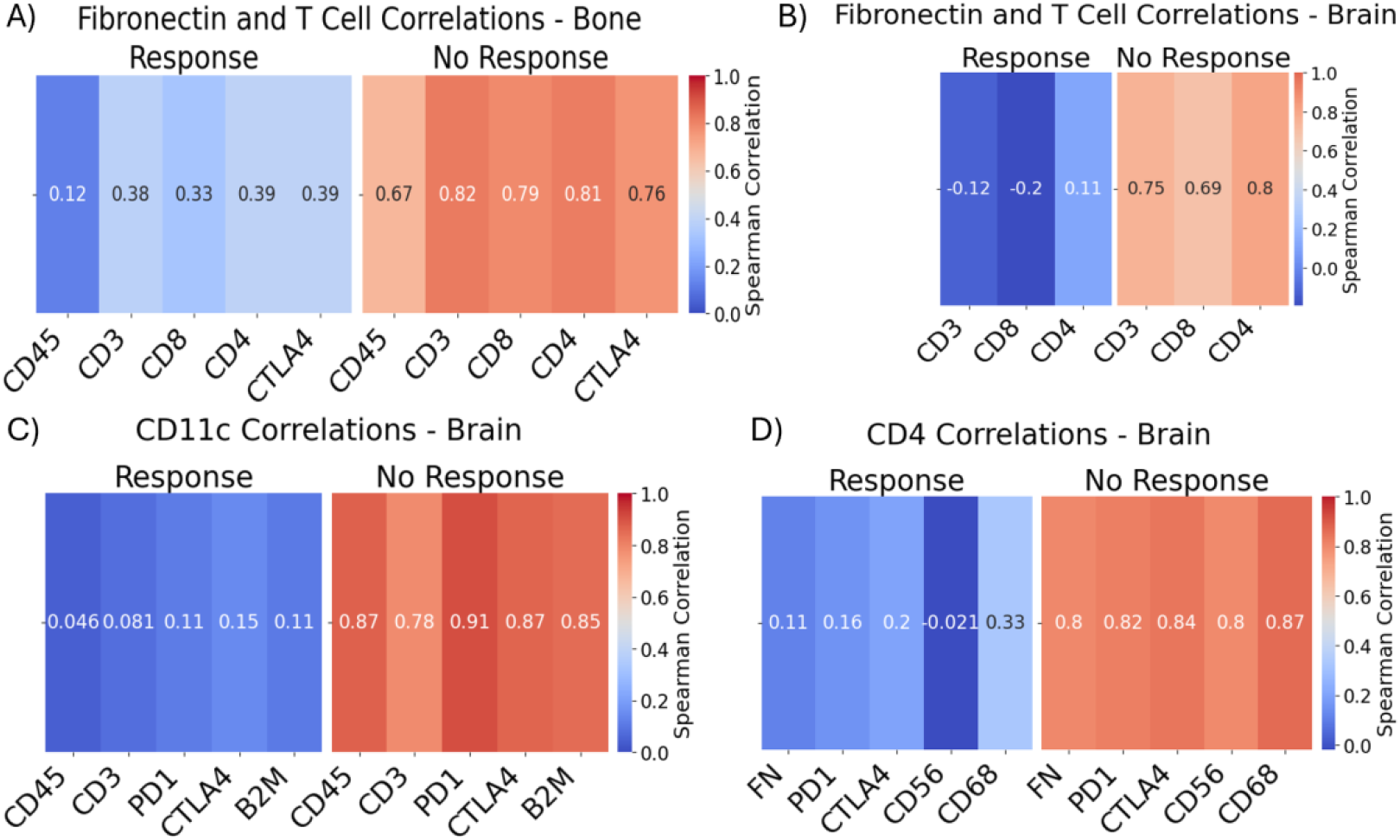
Fibronectin and immune marker correlations in bone and brain by response status. (A) Fibronectin correlations with T cell markers in bone. B) Fibronectin correlations with T cell markers in brain. C) CD11c correlations with other immune markers in brain. D) CD4 correlations with other immune markers in brain. FN = fibronectin. p < 0.05 for all R vs. NR based on Fisher r-to-z transformation.

CD11c correlations further support a coordinated immunosuppressive environment in brain NR samples (Fig. 9C). In these tissues, CD11c expression strongly correlated with markers of antigen presentation (B2M, ρ = 0.85), pan-leukocyte presence (CD45, ρ = 0.87), and key immune checkpoints (PD1, ρ = 0.91; CTLA4, ρ = 0.87). These associations were nearly absent in responders (ρ = 0.046–0.15), suggesting that in NR, dendritic cells sequester exhausted immune cells, facilitating an immunosuppressive niche that promotes tumor persistence, as previously described (Dähling et al., 2022).

Similarly, CD4 correlations revealed pronounced immune checkpoint engagement in NR (Fig. 9D). In the brain, CD4 expression was tightly correlated with PD1, CTLA4, CD68, and CD56 (ρ > 0.80 for all), suggesting that helper T cells are co-localized with exhausted and potentially suppressive immune populations, including myeloid cells and NK-like subsets. In contrast, these relationships were weak or absent in R tissues, indicating a more effective decoupling from suppressive immune interactions.

Together, these findings point to a shared immune-exclusion phenotype across bone and brain NR tissue samples, where T cells, myeloid cells, and immune checkpoints are spatially coordinated within fibrotic regions. By comparison, R tissues display a more disaggregated spatial immune architecture.

## Discussion

Despite the clinical success of T-DXd in HER2+ and HER2-low metastatic breast cancer, approximately 40–50% of patients fail to respond, underscoring a critical need to elucidate resistance mechanisms. This study utilized NanoString spatial proteomics platform to systematically profile the immune and stromal landscapes across multiple metastatic sites in T-DXd–responsive (R) versus –nonresponsive (NR) patients. We identify consistent and spatially resolved microenvironmental features that differentiate responders from NR, including HER2 heterogeneity, patterns of immune infiltration, and spatial coordination between stromal and immune compartments.

A central finding is that HER2 heterogeneity is significantly higher in NR, both at the whole-slide and ROI levels (Fig. 5). Interestingly, in R tissues, HER2 score negatively correlated with cytotoxic immune markers (e.g., CD8, GZMB), suggesting that in effective responders, cytotoxic cells may preferentially localize away from HER2-dense regions, possibly reflecting spatial immunoediting or tumor immune evasion. In contrast, NR samples displayed the opposite trend (Fig. 5C), indicating that even in the presence of target antigen and immune effector cells, antitumor activity remains blunted—highlighting the concept of immune dysfunction or exhaustion despite apparent immune presence.

Pan-cancer analysis revealed that fibronectin and GZMB were consistently upregulated in NR samples, regardless of prior treatment class (Fig. 6A–B, Table 5). While fibronectin likely contributes to immune exclusion through ECM remodeling, elevated GZMB expression—despite its association with cytotoxic lymphocytes—may reflect ineffective or exhausted immune activity, particularly in the context of high CD68+ macrophage presence. Notably, fibronectin was enriched specifically in tumor-sparse NR ROIs, suggesting that fibrotic barriers may obstruct immune access in regions where cancer cell density is low (Fig. 6E).

**Table 5:**
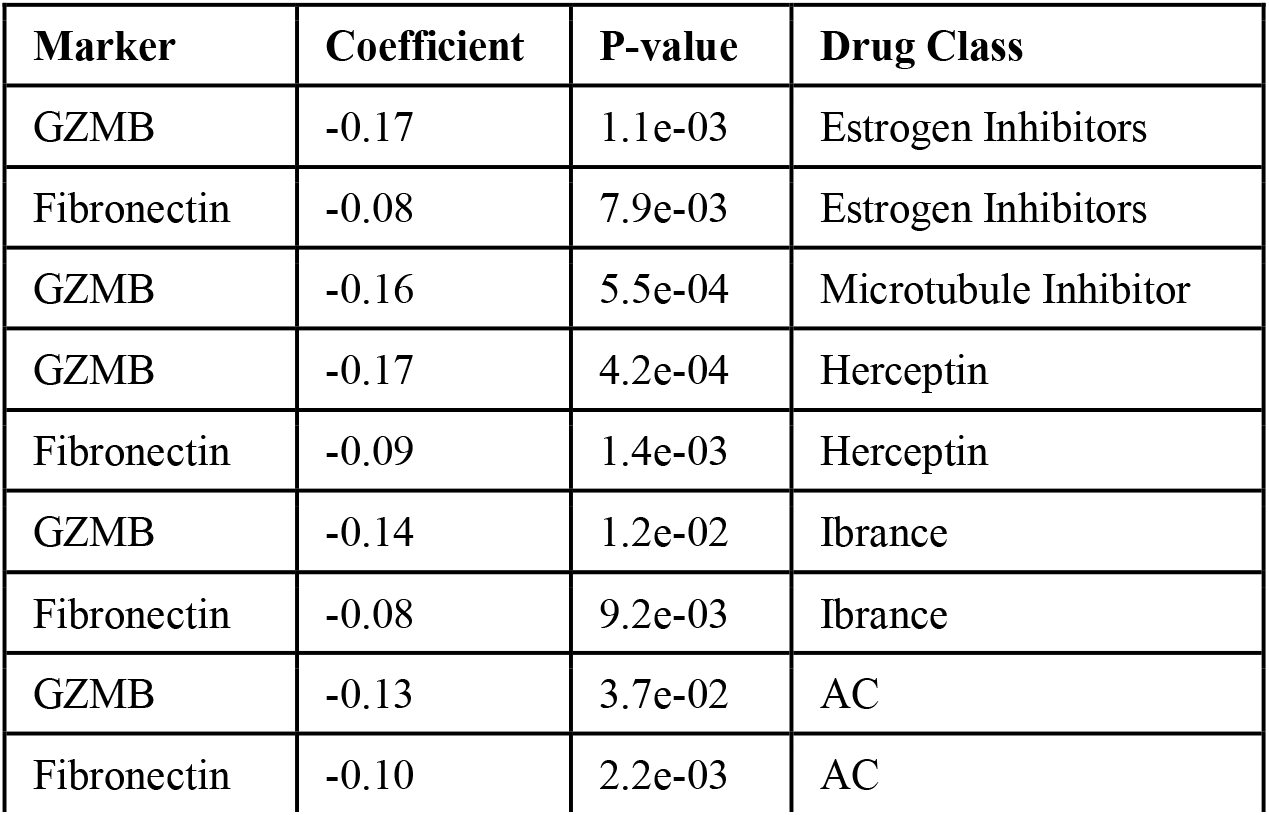
Linear mixed modeling coefficients for significant markers in predicting R vs. NR given treatment history.

Organ-specific analysis uncovered both conserved and divergent immune architectures. In bone and brain metastases, fibronectin showed strong positive correlations with CD3, CD4, and CD8 (ρ > 0.75–0.80) in non-responders, a pattern absent or reversed in responders (Fig 9A–B). These data suggest that in NR, fibronectin-rich niches trap immune cell populations, potentially sequestering them away from tumor cell contact or impairing their cytotoxic function. Moreover, in brain metastasis, CD11c and CD4 tightly correlated with immune checkpoints (PD1, CTLA4), B2M, and CD68 in NR (Fig 9C–D), suggesting spatially coordinated but functionally exhausted immune compartments dominated by myeloid cells. This immunosuppressive architecture may explain the poor efficacy of immune-targeted therapies in certain metastatic brain contexts. Future studies should investigate whether such correlations could also explain the poor immune engagement in HER2-high regions for NR tissues.

Importantly, this fibronectin–immune cell co-localization trend was reversed in lymph nodes (Fig. 8B). In NR lymph node metastases, fibronectin correlated negatively with immune markers such as CD45 (ρ = –0.24), CD56 (ρ = –0.12), and SMA (ρ = –0.10), whereas responders exhibited positive and coordinated interactions among ECM and immune markers (e.g., SMA– fibronectin ρ = 0.92). This striking reversal implies that fibronectin may have different functional roles across organ sites—serving as a physical barrier in bone and brain, while potentially guiding immune infiltration in lymph node metastases. Additionally, NR lymph nodes exhibited tight clustering of B and T cell markers (CD20–CD3/CD4/CD8; ρ > 0.85), possibly reflecting nonproductive lymphoid aggregation in the absence of effective immune activation.

These findings underscore the spatial complexity of tumor–immune–stroma interactions and point to fibronectin as a key mediator of immune architecture and function in mBC metastases. The observed organ-specific differences highlight the importance of considering tissue context in both biomarker interpretation and therapeutic design.

This study is limited by modest sample size. Nonetheless, the use of spatial proteomics enabled resolution of immunological heterogeneity at near-single-cell granularity. Future studies should incorporate spatial and single-cell transcriptomics and functional imaging to validate these immune phenotypes and explore combination therapies targeting the ECM or suppressive myeloid cells. Specifically, the opposing fibronectin–immune correlations between lymph nodes and other organs merit deeper mechanistic investigation, granularity.

In conclusion, our spatial proteomic analysis of HER2+ mBC metastases reveals that immune exclusion and ECM remodeling are key features of T-DXd resistance, site-specific biomarker interpretation is essential for mechanistic understanding, and site-specific immune architectures critically shape therapeutic outcomes. These insights provide a rationale for stratifying patients based on spatial immune phenotypes and for developing combination therapies that concurrently target ECM barriers and immunosuppressive myeloid populations to improve clinical response.

## Author Contributions

Conceptualization, H.Z., M.V., S.T.W., X.L.; data curation Y.G., W.Y., formal analysis, G.D.; investigation, G.D., J.Z., M.V.; methodology, M.V., H.Z, S.T.W., X.L.; resources, H.Z., S.T.W, X.L., Y.G., W.Y.; writing – original draft, G.D.; writing – review and editing, H.Z., S.T.W.; visualization, G.D.; supervision, H.Z., S.T.W., X.L.; project administration, H.Z., S.T.W., X.L.; funding acquisition, H.Z., S.T.W., X.L. All authors have read and agreed to the published version of the manuscript.

## Data Availability Statement

Data supporting reported results are contained in the figures and tables of the paper.

## Acknowledgments

We acknowledge the use of ChatGPT (OpenAI) for language refinement and grammar correction during manuscript preparation.

